# Environmental enrichment mitigates the long-lasting sequelae of perinatal fentanyl exposure

**DOI:** 10.1101/2021.07.31.454575

**Authors:** Jason Bondoc Alipio, Lace Marie Riggs, Madeline Plank, Asaf Keller

**Affiliations:** Department of Anatomy & Neurobiology, Program in Neuroscience, University of Maryland School of Medicine, Baltimore, MD 21201; Department of Psychiatry, Division of Translational and Basic Science, Program in Neuroscience and Training Program in Integrative Membrane Biology, University of Maryland School of Medicine, Baltimore, MD 21201

**Keywords:** Perinatal opioid exposure, fentanyl, environmental enrichment, plasticity, somatosensory cortex, anxiety, attention deficit hyperactivity disorder, adolescence, intervention

## Abstract

The opioid epidemic is a rapidly evolving societal issue driven, in part, by a surge in synthetic opioid use. A rise in fentanyl use among pregnant women has led to a 40-fold increase in the number of perinatally-exposed infants in the past decade. These children are more likely to develop mood- and somatosensory-related conditions later in life, suggesting that fentanyl may permanently alter neural development. Here, we examined the behavioral and synaptic consequences of perinatal fentanyl exposure in adolescent male and female C57BL/6J mice and assessed the therapeutic potential of environmental enrichment to mitigate these effects. Dams were given *ad libitum* access to fentanyl (10 µg/mL, *per os*) across pregnancy and until weaning (PD 21). Perinatally-exposed adolescent mice displayed hyperactivity (PD 45), enhanced sensitivity to anxiogenic environments (PD 46), and sensory maladaptation (PD 47) – sustained behavioral effects that were completely normalized by environmental enrichment (PD 21-45). Additionally, environmental enrichment normalized the fentanyl-induced changes in the frequency of miniature excitatory postsynaptic currents of layer 2/3 neurons in the primary somatosensory cortex (S1). We also demonstrate that fentanyl impairs short- and long-term potentiation in S1 layer 2/3 neurons which, instead, exhibit a sustained depression of synaptic transmission that is restored by environmental enrichment. On its own, environmental enrichment suppressed long-term depression of control S1 neurons from vehicle-treated mice subjected to standard housing conditions. These results demonstrate that the lasting effects of fentanyl can be ameliorated with a non-invasive intervention introduced during early development.

**Significance Statement:** Illicit use of fentanyl accounts for a large proportion of opioid-related overdose deaths. Children exposed to opioids during development have a higher risk of developing neuropsychiatric disorders later in life. Here, we employ a preclinical model of perinatal fentanyl exposure that recapitulates these long-term impairments and show, for the first time, that environmental enrichment can reverse deficits in somatosensory circuit function and behavior. These findings have the potential to directly inform and guide ongoing efforts to mitigate the consequences of perinatal opioid exposure.

## Introduction

The prevalence of opioid use disorder has steadily increased since the early 1990s, with women representing more than one-third of opioid users in the United States (Degenhardt et al., 2018). While opioid overdose deaths were initially driven by misuse of prescription opioids (like morphine) and their illicit counterparts (e.g., heroin), synthetic opioids, such as fentanyl, are now the main catalyst of overdose deaths (O’Donnell et al., 2017; Jannetto et al., 2019). Just among women 30-34 years of age, the rate of these overdose deaths have increased 35-fold in the last two decades (from 0.31 to 11 deaths per 100,000; VanHouten et al., 2019). Opioid use by pregnant women is a growing public health concern, with a 400% increase in the number of infants born to opioid-using mothers (from 1.5 to 6.5 per 1000 infants born; Haight et al., 2018). Opioid use during pregnancy increases the risk of miscarriage, premature birth, and stillbirth (Whiteman et al., 2014) whereas the infants who are born suffer from neonatal opioid withdrawal syndrome (NOWS; Winkelman et al., 2018). Often, NOWS is met with morphine, methadone, and buprenorphine treatment immediately after birth (Sutter et al., 2014), likely further compounding the developmental consequences of perinatal opioid exposure itself.

Longitudinal clinical studies indicate an increased risk of developing psychiatric conditions among children with a history of *in utero* and iatrogenic opioid exposure (Sherman et al., 2019). Specifically, these children have a higher incidence of anxiety (Cubas and Field, 1993), attention-deficit hyperactivity (Ornoy et al., 2001; Nygaard et al., 2016; Schwartz et al., 2021), autism spectrum (Sherman et al., 2019), and sensory processing disorders (Kivistö et al., 2014). Impairments in sensory processing is of particular interest because it is a common symptom in syndromes like attention-deficit hyperactivity and autism spectrum disorders (Ayres, 1964; Robertson and Baron-Cohen, 2017). While infants suffering from NOWS typically receive treatment for the acute symptoms of opioid withdrawal, there is virtually no treatment approaches specifically designed to address the other neurodevelopmental consequences that these children face across their lifetime.

Environmental enrichment in rodents can reverse early neurodevelopmental insults (Nithianantharajah and Hannan, 2006) and improve cognitive (Rijzingen et al., 1997; Passineau et al., 2001) and emotional functioning later in life (Brenes et al., 2009).This is due in part to the restoration of synaptic plasticity (Bayat et al., 2015) and neurogenesis (Gaulke et al., 2005), which are believed to re-establish neuronal complexity (Wei et al., 2021) via brain-derived neurotrophic factor signaling (Chen et al., 2005). While the synaptic benefits of environmental enrichment are often ascribed to its actions within corticolimbic circuitry, some of the most fundamental studies to establish its importance to synaptic efficacy took place within the somatosensory cortex (Smail et al., 2020). We previously showed that perinatal fentanyl exposure increases anxiety-like behavior and impairs synaptic transmission in anterior cingulate- and primary somatosensory (S1) cortex (Alipio et al., 2021a, 2021b). Because environmental enrichment can reduce anxiety-like behavior and restore the functional integrity of cortical circuits (Brenes et al., 2009; Wei et al., 2021), we hypothesized that environmental enrichment would mitigate the behavioral and synaptic deficits of perinatal fentanyl exposure.

## Materials and Methods

### Animals

All procedures were reviewed and approved by the University of Maryland Institutional Animal Care and Use Committee and adhered to the National Institutes of Health guide for the care and use of laboratory animals and Animal Research Reporting of In Vivo Experiments guidelines. Male and female C57BL/6J mice were used and bred in our temperature and humidity-controlled vivarium. When copulatory plugs were identified, we removed the sires and added fentanyl (10 µg/mL in 2% saccharin) or vehicle control (2% saccharin) to the water hydration pouches for *ad libitum* access by dams. Offspring were weaned at postnatal day (PD) 21 and housed 2-5 mice/cage/sex in standard housing, or 6-8 mice/cage/sex in enriched housing. Enriched housing was custom made from two One Cage 2100™ Micro-Isolator systems (Lab Products LLC, Seaford, DE). Enriched housing had a floor area of 420 in^2^ and included a variety of solid items for the mice to interact with (Fig 1. and Table 1). Food and water were available *ad libitum* and lights were maintained on a 12-hour cycle (0700 lights-on). Mice were tested in the open field test on PD 45 and in the tactile withdrawal test on PD 47. Slice experiments took place from PD 48-55.

**Figure 1.**
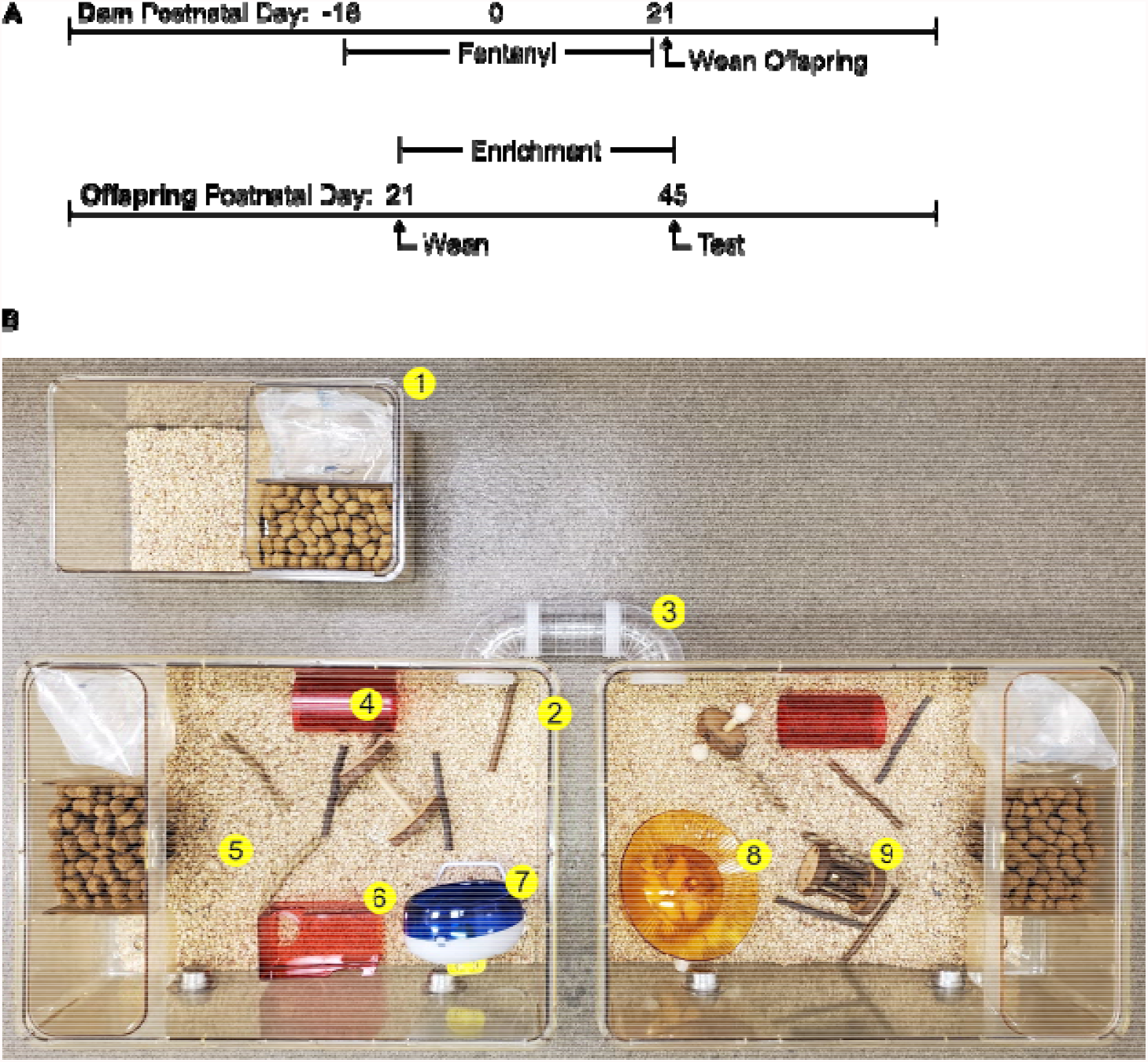
Experimental timeline of exposure to perinatal fentanyl and postnatal housing conditions. (**A**) Timeline depicting fentanyl exposure of mouse dams throughout pregnancy until weaning of offspring on postnatal day (PD) 21. (**B**) Offspring were weaned into standard (top cage) or enriched (bottom cage) housing conditions until behavioral (PD 45/47) and electrophysiological (PD 48-55) analyses took place. Enrichment items are numbered (1-9) and listed in Table 1.

**Table 1.** Product information for individual housing items

| Item | Description | Product Info | Company |
| --- | --- | --- | --- |
| 1 | Standard housing | Mouse 750<sup>TM</sup> ventilated cage | Lab Products |
| 2 | Environmental enrichment housing | One Cage 2100<sup>TM</sup> | Lab Products |
| 3 | Connecting tunnel | DIY clear tunnel | POPETPOP |
| 4 | Tunnel | Bio-Serv<sup>TM</sup> Mouse Tunnel | Fisher Scientific |
| 5 | 1" deep bedding | Corn Cob Bedding | Lab Supply |
| 6 | Platform | Bio-Serv<sup>TM</sup> Mouse Retreat | Fisher Scientific |
| 7 | Standing running wheel | Silent Spinner Wheel 4.5" | Kaytee |
| 8 | Dome and plate running wheel | Bio-Serv<sup>TM</sup> InnoDome and InnoWheel | Fisher Scientific |
| 9 | Variety chew toys | Small animal chew toys | Niteangel |

### Statistical Analyses

We adhered to accepted standards for rigorous study design and reporting to maximize the reproducibility and translational potential of our findings as described in Landis et al. (Landis et al., 2012) and in ARRIVE Guidelines (Kilkenny et al., 2010). Dams were randomly allocated to receive fentanyl or vehicle. In all our experiments, the primary end of points were prospectively defined for each experiment. All experimenters were blind to treatment conditions throughout data collection, scoring, and analysis. Statistical analyses were conducted using Prism v9 (GraphPad, San Diego, CA) and the minimum sample size was determined using G*Power Software Suite (Heinrich-Heine, Universität Düsseldorf). Statistical significance was defined as *p* < 0.05. No significant sex differences were observed for any of the experimental outcomes, thus, data from male and female mice were combined. There were no differences between mice of different litters within the same drug exposure group, therefore, individual mice served as a single sample count. For electrophysiology data, all neurons from a single mouse were averaged and served as a single sample count (Editorial, 2018). Student’s *t*-tests were used for two-group comparisons and the effect size was determined using Cohen’s *d* and defined as small (≤ 0.2), medium (≤ 0.5), or large (≥ 0.8). One-way repeated measures analysis of variance (AVOVA) was used when housing condition (standard vs. enriched) and time (repeated-measure) were independent variables. Two-way ANOVA was used when drug condition (vehicle vs. fentanyl) and housing condition were independent variables. Three-way repeated-measures ANOVA was used when drug condition, housing condition, and time were independent variables. When a significant main effect and/or interaction was observed, Tukey or Bonferroni *post-hoc* tests were used to assess pairwise comparisons and partial eta squared (Pη^2^) was used to determine if the effect size of the comparison was small (≤ 0.01), medium (≤ 0.06), or large (≥ 0.14). Fisher’s exact tests were used for contingency occurrence of plasticity (LTP vs. no LTP; LTD vs. no LTD) and *odds ratio* was used to determine if the effect size was small (≤ 1.49), medium (≤ 3.45), or large (≥ 9). Kruskal-Wallis tests were used for cumulative probability plots and Pη^2^ was used to determine effect size of those comparisons. Nonparametric alternatives were used if the data did not pass Spearman’s test for heteroscedasticity and D’Agostino-Pearson omnibus K2 test of distribution normality. For two-group nonparametric comparisons, Glass’ *delta* or Hedges’ *g* were used to determine if the effect size was small (≤ 0.2), medium (≤ 0.5), or large (≥ 0.8) whereas Pη^2^ was used to determine the effect size when three or more groups were being compared.

*Open Field Test* was used to assess general locomotor activity and anxiety-like behavior. Mice habituated to the testing room for at least 1-hr before the testing session began. Mice were individually placed in an open field arena (49 × 49 × 49 cm; San Diego Instruments, San Diego, CA) along the outside edge, facing the wall, and were allowed to freely explore the chamber for 30-min. The testing room was dimly lit with warm incandescent floor lamps and the center floor of the chamber read ∼30 lux. The test was recorded by an overhead digital camera, and distance traveled (cm) and time spent in the center (defined as 50% of the center area) were automatically scored using TopScan Software (CleverSys Inc, Reston, VA). Reduced time spent in the center zone of the arena is considered an anxiety-like response in this task.

*Tactile Hind Paw Withdrawal Test* was used to assess sensory threshold and adaptation. Mice habituated to the testing room for at least 1-hr before the testing session began, and were habituated to an elevated clear plexiglass box with a mesh bottom for 10 min. We applied von Frey filaments of increasing forces (in grams: 0.16, 0.40, 0.60, 1.00, 1.40, 2.00) to the plantar surface of the hind paw. A response was defined as an active withdrawal of the paw from the probing filament. The filament was applied to the same paw throughout the test. We used the up-down method to determine withdrawal threshold, as previously described (Dixon, 1965; Chaplan et al., 1994; Deuis et al., 2017). To assess sensory adaptation, we applied a von Frey filament one step above threshold to the plantar surface of the hind paw, opposite to the hind paw used during tactile sensitivity testing. The filament was applied once every 30 s until the animal stopped responding. The number of times the animals responded was counted, with persistent responding to tactile stimulation indicating sensory maladaptation

### Drugs and Solutions

Dams’ water hydration pouches contained either 10 µg/mL fentanyl citrate (calculated as free base) in 2% (w/v) saccharin or 2% saccharin (vehicle control), replenished weekly until litters were weaned on PD 21. Artificial cerebrospinal fluid (ACSF) compositions and slice preparations were based on slice collection methods of Ting et al. (Ting et al., 2014) *N*-methyl-D-glucamine (NMDG) ACSF contained (in mM): 92 NMDG, 30 NaHCO_3_, 20 HEPES, 25 glucose, 5 Na-ascorbate, 2 thiourea, 1.25 NaH_2_PO_4_, 2.5 KCl, 3 Na-pyruvate, 0.5 CaCl_2_·2H_2_O and 10 MgSO_4_·7H_2_O. Holding ACSF contained (in mM): 120 NaCl, 2.5 KCl, 1.25 NaH_2_PO_4_, 24 NaHCO_3_, 12.5 glucose, 2 MgSO_4_·7H_2_O, and 2 CaCl_2_·2H_2_O. Recording ACSF contained (in mM): 120 NaCl, 3 KCl, 1 NaH_2_PO_4_, 25 NaHCO_3_, 20 glucose, 1.5 MgSO_4_·7H_2_O, and 2.5 CaCl_2_·2H_2_O. ACSF pH was adjusted to 7.4 and osmolarity to 305 ± 2 mOSm. All ACSF solutions were saturated with carbogen (95% O_2_ and 95% CO_2_). Patch pipettes contained (in mM): 130 cesium methanesulfonate, 10 HEPES, 1 magnesium chloride, 2.5 ATP-Mg, 0.5 EGTA, 0.2 GTP-Tris, 5 QX-314, and 2% biocytin. The pH of the internal pipette solution was adjusted to 7.3 and osmolarity to 290 ± 2 mOSm. To isolate excitatory postsynaptic currents (EPSCs), gabazine (1 µM) was included in the ACSF. For miniature EPSC (mEPSC) recordings, tetrodotoxin (1 µM) was included in the ACSF. All recordings were obtained at room temperature.

### Slice Preparation

Mice were deeply anesthetized with intraperitoneal injection of ketamine (180 mg/kg) and xylazine (20 mg/kg) then transcardially perfused with ice-cold (4 °C) NMDG ACSF. Their brains were rapidly extracted following decapitation. Coronal slices (300 µm thick) containing the primary somatosensory cortex (S1) were cut in ice-cold (4 °C) NMDG ACSF using a Leica VT1200s vibratome (Leica Biosystems, Buffalo Grove, IL) and transferred to warm (33 °C) NMDG ACSF for 10 min. The slices were then transferred to warm (33 °C) holding ACSF and allowed to cool to room temperature (20-22 °C) for at least 45 min before electrophysiology recordings.

### Electrophysiology

Whole cell patch-clamp recordings were obtained from S1 layer 2/3 neurons with a Multiclamp 700B amplifier (Molecular Devices, San Jose, CA) low-pass filtered at 1.8 kHz with a four-pole Bessel filter, and digitized with Digidata 1440A (Molecular Devices). Slices were placed in a submersion chamber and continually perfused (>2 mL/min) with recording ACSF. Neurons were visually identified by infrared differential interference contrast imaging and location and neuronal morphology verified after each recording with biocytin immunohistochemistry. Borosilicate patch pipettes had an impedance of 4-6 MΩ. Once GΩ seal was obtained, neurons were held in voltage-clamp configuration at -70 mV and the input resistance, resting membrane potential, and capacitance were measured. Series resistance (<30 MΩ) was monitored throughout recordings and recordings were discarded if series resistance changed by >20% from baseline. Concentric bipolar tungsten electrodes were used to deliver electrical stimulation (0.2 ms duration) in S1 layer 5, below the recorded neuron. Electrically-evoked current responses were recorded at 1.5-fold threshold, defined as the minimum stimulation intensity required to produce a visible current response beyond baseline noise.

Long-term potentiation (LTP) was induced using a pairing induction protocol: 80 stimulation pulses at 2 Hz paired with postsynaptic depolarization to +30 mV (Zhao et al., 2005; Toyoda et al., 2009). Long-term depression (LTD) was induced using low frequency stimulation: 900 stimulation pulses at 1 Hz. The occurrence of LTP or LTD was assessed by comparing the average EPSC amplitude from 5-10 min of the baseline to 25-30 min after the plasticity induction protocol (Nicoll, 2017). EPSC responses to electrically evoked paired pulse stimulation (50 ms intervals) were recorded after the 10 min baseline and again 30 min after the plasticity induction protocol. The paired pulse ratio (PPR) was obtained by averaging the mean amplitude of the second EPSC by that of the first (Kim and Alger, 2001). We recorded and analyzed 3 min long segments of mEPSCs in separate slices that did not undergo electrical stimulation. Autodetection parameters for inclusion of events was determined by calculating minimum threshold: root mean square (RMS)^2^ × 1.5. To control for oversampling and unequal sample size between groups, we performed quantile sampling of mEPSC frequency and amplitude by computing 29 evenly spaced quantile values from each neuron, starting at the 1^st^ percentile and ending at the 94.2^th^ percentile, with a step size of 3.33% (Hanes et al., 2020). Data acquisition was performed using Clampex and analyzed with Clampfit (Molecular Devices). Mini Analysis software (Synaptosoft, USA) was used to analyze mEPSC recordings.

## Results

### Perinatal fentanyl exposure

We administered fentanyl citrate (10 µg/mL) in the drinking water of pregnant mouse dams throughout their pregnancy and until their litters were weaned at postnatal day (PD) 21 (Fig. 1A). We chose 10 µg/ml since it is the optimal concentration mice will readily self-administer without producing motor deficits (Wade et al., 2008, 2013) and is well below the mouse oral LD50 (fentanyl citrate MSDS, Cayman Chemical, Ann Arbor, MI). We have recently found that perinatal treatment with 10 µg/ml results in behavioral and synaptic deficits in adolescent mice (Alipio et al., 2021b). Offspring were weaned into either standard or enriched housing environments (Fig. 1, Table 1).

### Environmental enrichment reverses the sustained behavioral deficits that are induced by perinatal fentanyl exposure

#### Hyperactivity

We used the open field test (Fig. 2A) to examine general locomotor behavior of adolescent male and female mice perinatally exposed to either vehicle or fentanyl, raised in either standard or an enriched housing environment (Fig. 2B-C; *n* = 15-20 mice per group, about 50/50 male to female ratio). There was no significant sex × drug × housing interaction (*F* (1, 59) = 2.83, *p* = 0.09) nor any differences between sex within each drug and housing group (Tukey’s post hoc, *p* > 0.05) on total distance traveled. Therefore, data from both sexes were grouped according to drug and housing condition. All mice habituated to the open field over time, in that distance traveled progressively decreased over the course of the procedure (Fig. 2B; *F* (3.7, 233.7) = 119.70, *p* < 10^−4^, Pη^2^ = 0.65, *large effect*). There was a significant time × drug (Three-way RM ANOVA, *F* (5, 310) = 6.69, *p* < 10^−4^, Pη^2^ = 0.09, *medium effect*) and drug × housing interaction (*F* (1, 62) = 7.07, *p* = 0.009, Pη^2^ = 0.10, *medium effect*). There was a significant drug × housing interaction in total distance traveled (Fig. 2C; Two-way ANOVA, *F* (1, 62) = 7.07, *p* = 0.01, Pη^2^ = 0.10, *medium effect*), with fentanyl-exposed, standard-housed mice exhibiting higher total distance traveled than vehicle-exposed, standard-housed mice (Tukey’s post hoc, *p* = 0.01, Cohen’s *d* = 1.23, *large effect*). While environmental enrichment did not influence locomotor behavior on its own (i.e., in vehicle-exposed mice; Tukey’s post hoc, *p* = 0.97), it attenuated the hyperactivity of mice perinatally exposed to fentanyl (Tukey’s post hoc, *p* = 0.007, Cohen’s *d* = 1.58, *large effect*). These data suggest that perinatal fentanyl exposure leads to hyperactivity and that environmental enrichment can reverse this effect without itself perturbing locomotor behavior.

**Figure 2.**
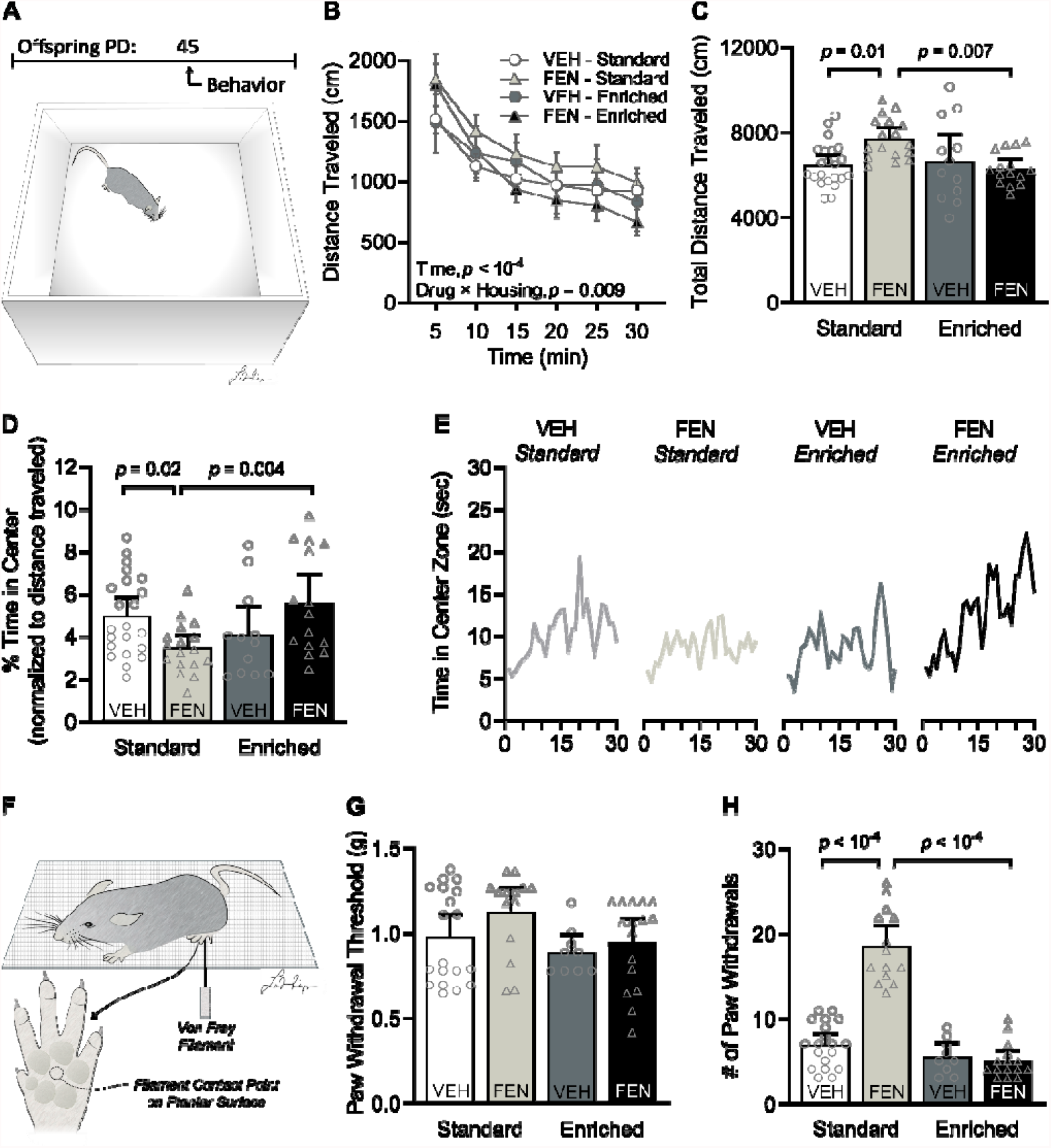
Environmental enrichment reverses the sustained behavioral deficits that are induced by perinatal fentanyl exposure. (**A**) Timeline depicting behavioral assays and a schematic of the open field test. *n* = 15-20 mice per group. (**B, C**) Perinatal fentanyl exposure increases distance traveled across time. (**D, E**) Perinatal fentanyl exposed mice raised with environmental enrichment have comparable distance traveled with vehicle controls but spend less time in the center area of the open field. Fentanyl-exposed mice raised with environmental enrichment spend more time in the center area of the open field, comparable to vehicle controls. (**F**) Graphic depicting von Frey tactile stimulation on the plantar surface of the hind paw. (**G**) There were no differences across groups in the paw withdrawal threshold to tactile stimulation. *n* = 11-19 mice per group. (**H**) Standard housed fentanyl exposed mice have a higher number of paw withdrawal responses to repeated stimuli than standard housed vehicle mice. Fentanyl exposed mice raised with environmental enrichment have a lower number of paw withdrawal responses, comparable to vehicle mice raised in standard and enriched environment. Data depict means, *p* values, and 95% confidence intervals.

### Anxiety-like behavior

To test our prediction that environmental enrichment will attenuate anxiety-like behavior, we assessed time spent in the center zone during the open field test (Fig. 2D; *n* = 15-20 mice per group; Kruskal-Wallis test, *H* = 10.17, *p* = 0.01, Pη^2^ = 0.01, *small effect*). Fentanyl-exposed, standard-housed mice spent less time in the center zone than did vehicle-exposed, standard-housed mice (Dunn’s post hoc, *p* = 0.02, Glass’ *delta* = 0.79, *medium effect*), and this, too, was selectively reversed by environmental enrichment (Dunn’s post hoc, *p* = 0.004, Glass’ *delta* = 0.88, *large effect*). The observation that enrichment improves center-zone exploration of fentanyl-exposed mice is notable, given the sustained hyperlocomotor/anxiogenic effect induced by fentanyl under standard-housing conditions. This is further evidenced by fentanyl-exposed, standard-housed mice spending less time in the center zone over the course of the procedure relative to their enriched-housed fentanyl-exposed counterparts (Fig. 2E; Kruskal-Wallis test, *H* = 14.80, *p* = 0.002, Pη^2^ = 0.01, *small effect*; Dunn’s post hoc, *p* = 0.01, Glass’ *delta* = 1.15, *large effect*), with the hyperlocomotor phenotype promoting center zone entries among fentanyl-exposed, standard-housed. These data suggest that environmental enrichment reduces anxiety-like behavior in adolescent mice that were perinatally exposed to fentanyl.

### Sensory adaptation

We tested sensory adaptation, a reduction in sensitivity to repeated stimuli, by applying von Frey filaments to the plantar surface of the hind paw (Fig. 2F-H; *n* = 11-19 mice per group). To test whether mice can sense tactile stimuli, we assessed hind paw withdrawal to threshold stimulation (Fig. 2G). There was no interaction nor main effect of drug and housing condition on paw withdraw threshold, suggesting that all mice similarly perceive tactile stimuli (Kruskal-Wallis test, *H* = 6.80, *p* = 0.07). Next, we tested whether mice adapt to repeated application of von Frey filaments above their threshold hind paw withdrawal response (Fig. 2H). There was a significant drug × housing interaction (Two-way ANOVA, *F* (1, 54) = 56.27, *p* < 10^−4^, Pη^2^ = 0.51, *large effect*). Fentanyl-exposed, standard-housed mice continued to respond to the stimuli more than twice as much as vehicle-exposed, standard-housed mice (Tukey’s post hoc, *p* < 10^−4^, Glass’ delta = 4.31, *large effect*). Enriched housing completely reversed this sensory maladaptation (Tukey’s post hoc, *p* < 10^−4^, Glass’ delta = 6.22, *large effect*). These data suggest that perinatal fentanyl exposure leads to a sustained impairment in tactile sensory adaptation that can be reversed by environmental enrichment. Notably, while perinatal fentanyl exposure promotes hyperactivity and anxiety-like behavior relative to vehicle-exposed mice with large and medium effect, respectively, its influence on sensory adaptation was larger. These data suggest that fentanyl leads to an enduring sequalae of behavioral changes that can be fully reversed by environmental enrichment, with sensory maladaptation possibly being among the most prominent changes induced by perinatal opioid exposure.

### Environmental enrichment restores long-term potentiation in S1 layer 2/3 neurons after perinatal fentanyl exposure

The fentanyl-induced impairment in sensory adaptation, as well as its reversal by environmental enrichment, suggests that these changes may be associated with altered synaptic plasticity in somatosensory cortex (S1). To test this, we assessed whether perinatal fentanyl exposure and environmental enrichment influenced synaptic potentiation of excitatory postsynaptic currents (EPSCs) in S1 layer 2/3 neurons (Fig. 3; *n* = 6-8 mice/group, 1 neuron/mouse, about 50/50 male to female ratio). There was no significant sex × drug × housing interaction (*F* (1, 21) = 1.21, *p* = 0.28) nor any differences between sex within each drug and housing group (Tukey’s post hoc, *p* > 0.05) on long-term plasticity. Therefore, data from both sexes were grouped according to drug and housing condition.

**Figure 3.**
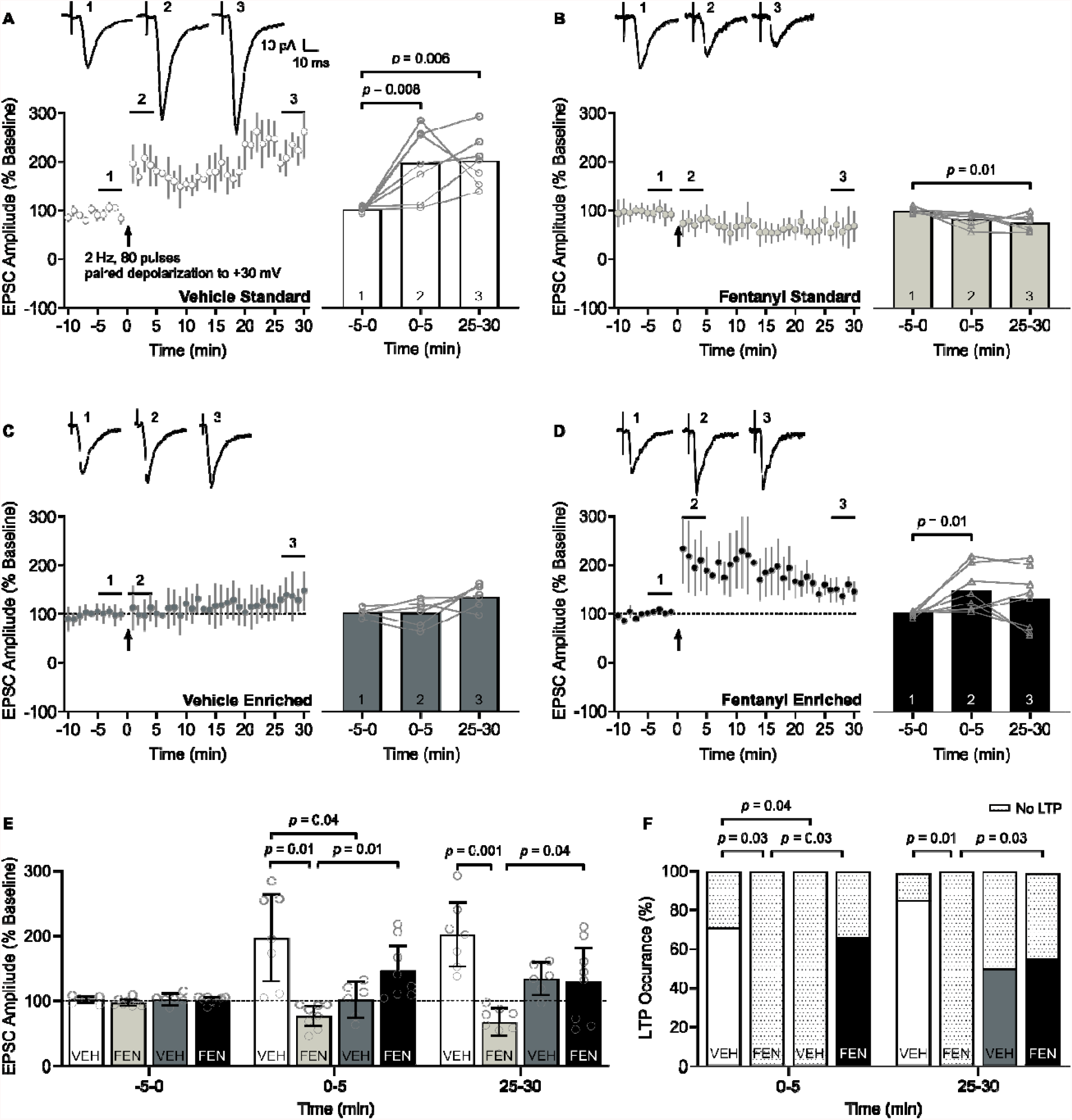
Environmental enrichment restores potentiation in S1 layer 2/3 neurons after perinatal fentanyl exposure. (**A-D**) Time course (left) of short- and long-term potentiation (STP/LTP) of excitatory postsynaptic currents (EPSCs) following paired induction (80 electrical stimulation pulses at 2 Hz paired with postsynaptic depolarization to +30 mV). Bar graphs (right) show group data comparing EPSC amplitudes at baseline -5-0 min (1), STP 0-5 min (2), and LTP 25-30 min (3) after LTP induction. Insets depict sample traces from times indicated on time course graph. (**A**) Within-group comparisons indicate STP and LTP were induced in S1 layer 2/3 neurons from vehicle-exposed, standard-housed mice. (**B**) LTP paired induction parameters induced long-term depression (LTD) of the EPSC in fentanyl-exposed, standard-housed mice. (**C**) LTP paired induction protocol failed to further potentiate EPSC amplitude from vehicle-exposed, enriched-housed mice. (**D**) The paired induction protocol induced STP, but not LTP, in fentanyl-exposed, enriched-housed mice. (**E**) Between-group comparisons of the EPSC amplitudes post LTP induction. Fentanyl-exposed, standard-housed mice had lower EPSC amplitudes than vehicle-exposed, standard-housed mice at 0-5 and 25-30 min timepoints. Environmental enrichment increased the EPSC amplitude of fentanyl-exposed, enriched-housed mice, relative to their fentanyl-exposed, standard-housed counterparts also at 0-5 and 25-30 min timepoints. (**F**) Between-group comparisons indicate fentanyl-exposed, standard-housed mice had a lower occurrence of neurons that exhibit STP and LTP than vehicle-exposed, standard-housed mice. Raising fentanyl-exposed mice in environmental enrichment increased the proportion of neurons that potentiate. *n* = 6-8 mice/group, 1 neuron/mouse. Data depict means and 95% confidence intervals.

Mechanisms that govern the expression of short-term potentiation (STP) and long-term potentiation (LTP) differ (Schulz and Fitzgibbons, 1997), and both may be involved in sensory adaptation behavior (Monday et al., 2018; Motanis et al., 2018), therefore we assessed both forms of synaptic potentiation among our groups. Both STP and LTP were readily induced in S1 layer 2/3 neurons from vehicle-exposed, standard-housed mice (Fig. 3A). Amplitudes of evoked EPSCs were higher at both STP 0-5 min (Bonferroni’s post hoc, *p* = 0.008, Cohen’s *d* = 1.86, *large effect*) and LTP 25-30 min (Bonferroni’s post hoc, *p* = 0.006, Cohen’s *d* = 2.63, *large effect*) timepoints after the LTP paired induction protocol (80 electrical stimulation pulses at 2 Hz paired with postsynaptic depolarization to +30 mV) relative to baseline -5-0 min (One-way RM ANOVA, *F* (2, 12) = 8.71, *p* = 0.004, Pη^2^ = 0.59, *large effect*). In contrast, neurons from fentanyl-exposed, standard-housed mice did not potentiate, and instead showed a long-term depression (LTD)-like effect with significantly reduced EPSC amplitudes at the 25-30 min timepoint (Bonferroni’s post hoc, *p* = 0.01, Cohen’s *d* = 1.92, *large effect*) relative to baseline (Fig. 3B; One-way RM ANOVA, *F* (2, 12) = 5.47, *p* = 0.02, Pη^2^ = 0.48, *large effect*). In vehicle-exposed mice raised with environmental enrichment, the induction parameters tested here failed to evoke a statistically significant change in EPSC amplitude from baseline (Fig. 3C; One-way RM ANOVA, *F* (2, 10) = 4.38, *p* = 0.04, Pη^2^ = 0.46, *large effect*; Bonferroni’s post hoc, *p* > 0.05). In fentanyl-exposed mice raised with environmental enrichment, the LTP-induction protocol produced STP at the 0-5 min timepoint (Bonferroni’s post hoc, *p* = 0.04, Cohen’s *d* = 1.42, *large effect*), but EPSC amplitude was not significantly different from baseline by 25-30 min post-induction (Fig. 3D; One-way RM ANOVA, *F* (2, 14) = 3.27, *p* = 0.06).

Consistent with these within-group comparisons, between-group comparisons revealed a complete block of STP and LTP in neurons from fentanyl-exposed, standard-housed mice and a restoration of STP and LTP in fentanyl-exposed mice by environmental enrichment, compared to their respective controls (Fig. 3E). There was a significant time × drug × housing interaction (Three-way RM ANOVA, *F* (2, 50) =11.43, *p* < 10^−4^, Pη^2^ = 0.31, *large effect*). At the 0-5 min timepoint, neurons from vehicle-exposed, enriched-housed mice had lower post-LTP EPSC amplitudes (Tukey’s post hoc, *p* < 10^−3^, Cohen’s *d* = 1.75, *large effect*) than their vehicle-exposed, standard-housed counterparts. Neurons from fentanyl-exposed, standard-housed mice had lower EPSC amplitudes than vehicle-exposed, standard-housed mice (Tukey’s post hoc, *p* < 10^−4^, Cohen’s *d* = 2.28, *large effect*). This reduction in fentanyl-exposed mice was restored by environmental enrichment (Tukey’s post hoc, *p* = 0.01, Cohen’s *d* = 2.02, *large effect*). At the LTP 25-30 min timepoint, neurons from fentanyl-exposed, standard-housed mice had lower EPSC amplitudes than vehicle-exposed, standard-housed mice (Tukey’s post hoc, *p* < 10^−4^, Cohen’s *d* = 3.22, *large effect*). This reduction was also restored by environmental enrichment (Tukey’s post hoc, *p* = 0.04, Cohen’s *d* = 1.33, *large effect*).

While nearly all neurons from vehicle-exposed, standard-housed mice exhibited STP (Fig. 3F; 5/7 neurons, 71.4%) and LTP (6/7 neurons, 85.7%), none of the neurons from fentanyl-exposed, standard-housed mice exhibited STP or LTP (0/7 neurons, 0%). Neurons from vehicle-exposed, enriched-housed mice did not exhibit STP (0/6 neurons, 0%), however, half of the neurons exhibited LTP (3/6 neurons, 50%). Neurons from fentanyl-exposed enriched-housed mice exhibited both STP and LTP (5/8 neurons, 62.5%). Between-group comparisons further indicate that fentanyl-exposed, standard-housed mice had a lower occurrence of both STP and LTP than vehicle-exposed, standard-housed mice (Fisher’s exact test, STP, *p* = 0.03, *odds ratio* = 33, *large effect*; LTP, *p* = 0.01, *odds ratio* = 65, *large effect*) and that environmental enrichment increased the proportion of neurons that exhibit STP and LTP in fentanyl-exposed mice compared to fentanyl-exposed, standard-housed mice (Fisher’s exact test, STP, *p* = 0.02, *odds ratio* = 23.57, *large effect*; LTP *p* = 0.02, *odds ratio* =23.57, *large effect*). These data demonstrate that perinatal fentanyl exposure impairs both STP and LTP in S1 layer 2/3 neurons from mice raised under standard housing conditions, and that environmental enrichment can restore potentiation to levels comparable to vehicle-exposed, standard-housed mice. Neurons from control mice raised in an enriched environment exhibit an attenuated and lower occurrence of STP and LTP relative to standard housing conditions, which may suggest an occlusive effect of enrichment on synaptic plasticity.

### Environmental enrichment suppresses long-term depression

We assessed whether perinatal fentanyl exposure influenced LTD of EPSCs in S1 layer 2/3 neurons and the impact of environmental enrichment on these changes (Fig. 4; *n* = 5-6 mice/group, 1 neuron/mouse). LTD of EPSC amplitudes were evident following low-frequency electrical stimulation (900 stimuli, 1 Hz) in S1 layer 2/3 neurons from vehicle-exposed, standard-housed mice (Fig. 4A; Paired *t*-test, *p* < 10^−4^, Cohen’s *d* = 11.13, *large effect*) and in fentanyl-exposed, standard-housed mice (Fig. 4B; Wilcoxon matched-pairs signed rank test, *p* = 0.03, Glass’ delta = 40.11, *large effect*). However, LTD was not induced in neurons from vehicle-exposed mice raised in an enriched environment (Fig. 4C; Wilcoxon matched-pairs signed rank test, *p* = 0.12). In contrast, LTD was readily induced in fentanyl-exposed, enriched-housed mice (Fig. 4D; Wilcoxon matched-pairs signed rank test, *p* = 0.03, Glass’ delta = 6.46, *large effect*).

**Figure 4.**
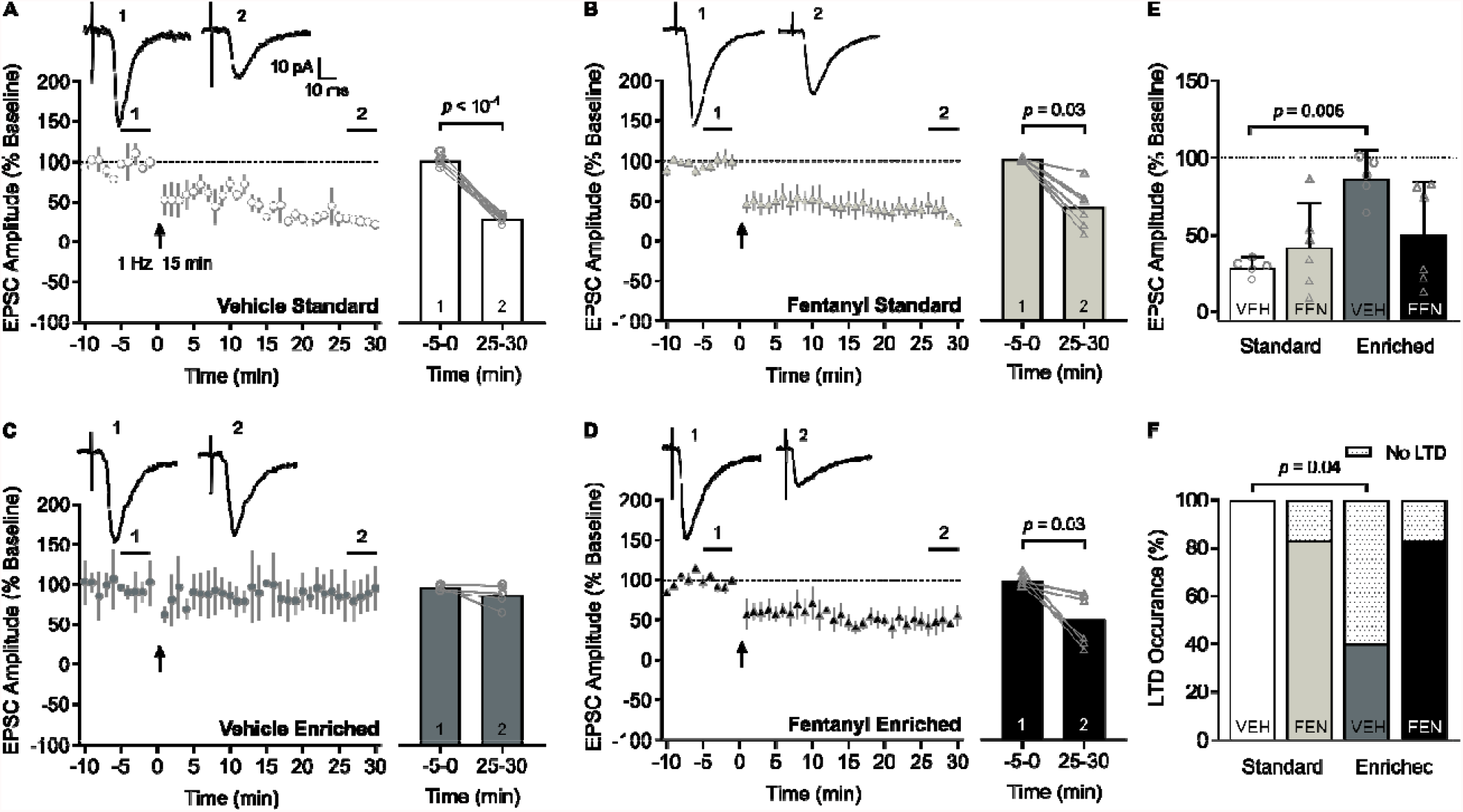
Environmental enrichment suppresses long-term depression. (**A-D**) Time course (left) of long-term depression (LTD) of excitatory postsynaptic currents (EPSCs) following low frequency stimulation (900 electrical stimulation pulses at 1 Hz). Bar graphs (right) show group data comparing EPSC amplitudes at baseline (1) and at 25-30 min after LTD induction (2). Insets depict sample traces from times indicated on time course graph. (**A**) Within-group comparisons indicate LTD was induced in S1 layer 2/3 neurons from vehicle-exposed, standard-housed mice and in (**B**) fentanyl-exposed, standard-housed mice. (**C**) LTD was not induced in vehicle-exposed, enriched-housed mice, but was in (**D**) fentanyl-exposed, enriched-housed mice. (**E**) Between-group comparisons of the EPSC amplitudes post-LTD induction indicate that vehicle-exposed, enriched-housed mice failed to induce LTD compared to their vehicle-exposed, standard-housed counterparts. (**F**) Between-group comparisons of the occurrence of LTD indicate vehicle-exposed, enriched-housed mice had a lower occurrence of neurons that exhibit LTD than vehicle-exposed, standard-housed mice. *n* = 5-6 mice/group, 1 neuron/mouse. Data depict means and 95% confidence intervals.

Consistent with this, between-group comparisons of post-LTD EPSC amplitudes revealed a significant drug × housing interaction (Fig. 4E; Two-way ANOVA, *F* (1, 18) = 5.86, *p* = 0.02, Pη^2^ = 0.24, *large effect*) with greater LTD in neurons from vehicle-exposed, standard-housed mice, relative to vehicle-exposed, enriched-housed mice (Tukey’s post hoc, *p* = 0.006, Cohen’s *d* = 5.26, *large effect*). All neurons from vehicle-exposed standard-housed mice exhibited LTD (5/5 neurons) whereas only 20% of neurons from vehicle-exposed, enriched-housed mice exhibited LTD (Fig. 4F; 1/5 neurons; Fisher’s exact test, *p* = 0.04, *odds ratio* = 40, *large effect*). Most neurons from fentanyl-exposed mice exhibited LTD (5/6 neurons 83.33%) in either housing condition, respectively.

These data demonstrate that perinatal fentanyl exposure does not appear to affect LTD induction or expression, regardless of housing conditions. Similar to LTP, LTD is readily induced in S1 layer 2/3 neurons of mice raised in standard, conventional cages, but not in neurons from mice raised in an enriched environment.

### Evoked glutamate release probability is not influenced by perinatal fentanyl exposure or environmental enrichment

We assessed paired pulse ratios (PPRs) to determine if the changes in plasticity induced by perinatal fentanyl exposure and environmental enrichment were mediated by presynaptic changes in vesicle release probability (Fig. 5). Figure 5A depicts sample traces from each of the experimental groups. Before LTP/LTD induction, there were no differences in baseline PPR among the experimental groups (Fig. 5B; *n* = 9-12 mice/group, 1 neuron/mouse; Kruskal-Wallis test, *p* > 0.05), nor did PPR change following the induction of LTP (Fig. 5C: *n* = 4-6 mice/group, 1 neuron/mouse; Paired *t*-test, *p* > 0.05) or LTD (Fig. 5D: *n* = 4-6 mice/group, 1 neuron/mouse; Paired *t*-test, *p* > 0.05). These data suggest that neither perinatal fentanyl exposure nor environmental enrichment influence the probability of evoked glutamate release at baseline or following induction of LTP or LTD in S1 layer 2/3 neurons with the induction protocols used in the current study.

**Figure 5.**
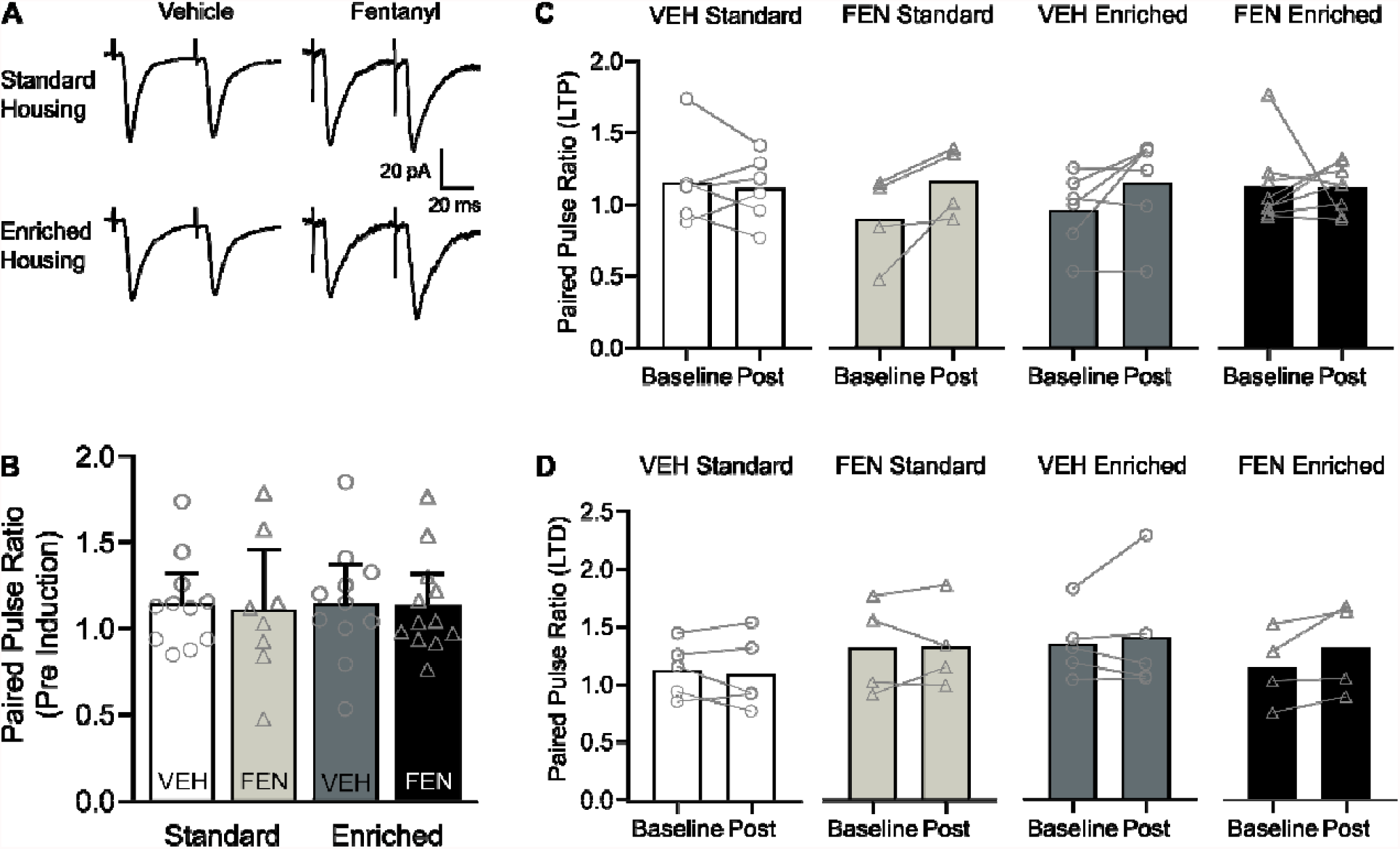
Evoked glutamate release probability is not influence by perinatal fentanyl exposure or environmental enrichment. (**A**) Representative traces of EPSC responses to paired pulse stimulation. (**B**) There were no differences in the paired pulse ratio (PPR) between groups prior to plasticity induction (*n* = 9-12 mice/group, 1 neuron/mouse). (**C, D**) Within each group, there were no differences in baseline PPR compared to PPR post-LTP or LTD induction (*n* = 4-6 mice/group, 1 neuron/mouse). Data depict means and 95% confidence intervals.

### Environmental enrichment restores changes in mEPSC frequency induced by perinatal fentanyl exposure

Some of the mechanisms and functions of spontaneous synaptic transmission are distinct from that of action potential-evoked synaptic transmission (Kavalali, 2015). To determine if perinatal fentanyl exposure influenced spontaneous excitatory synaptic transmission, we recorded miniature excitatory postsynaptic currents (mEPSCs) in S1 layer 2/3 neurons (Fig. 6; *n* = 6-8 mice/group, 1-2 neurons/mouse). We averaged all events from each neuron and animal for between-group comparisons of mEPSC frequency (Fig 6B; Kruskal-Wallis test, *H* = 12.41, *p* = 0.006, P η ^2^ = 0.03, *medium effect*). Neurons from vehicle-exposed, enriched-housed mice had higher mEPSC frequency than vehicle-exposed, standard-housed mice (Dunn’s post hoc, *p* = 0.01, Glass’ *delta* = 21.00, *large effect*).

**Figure 6.**
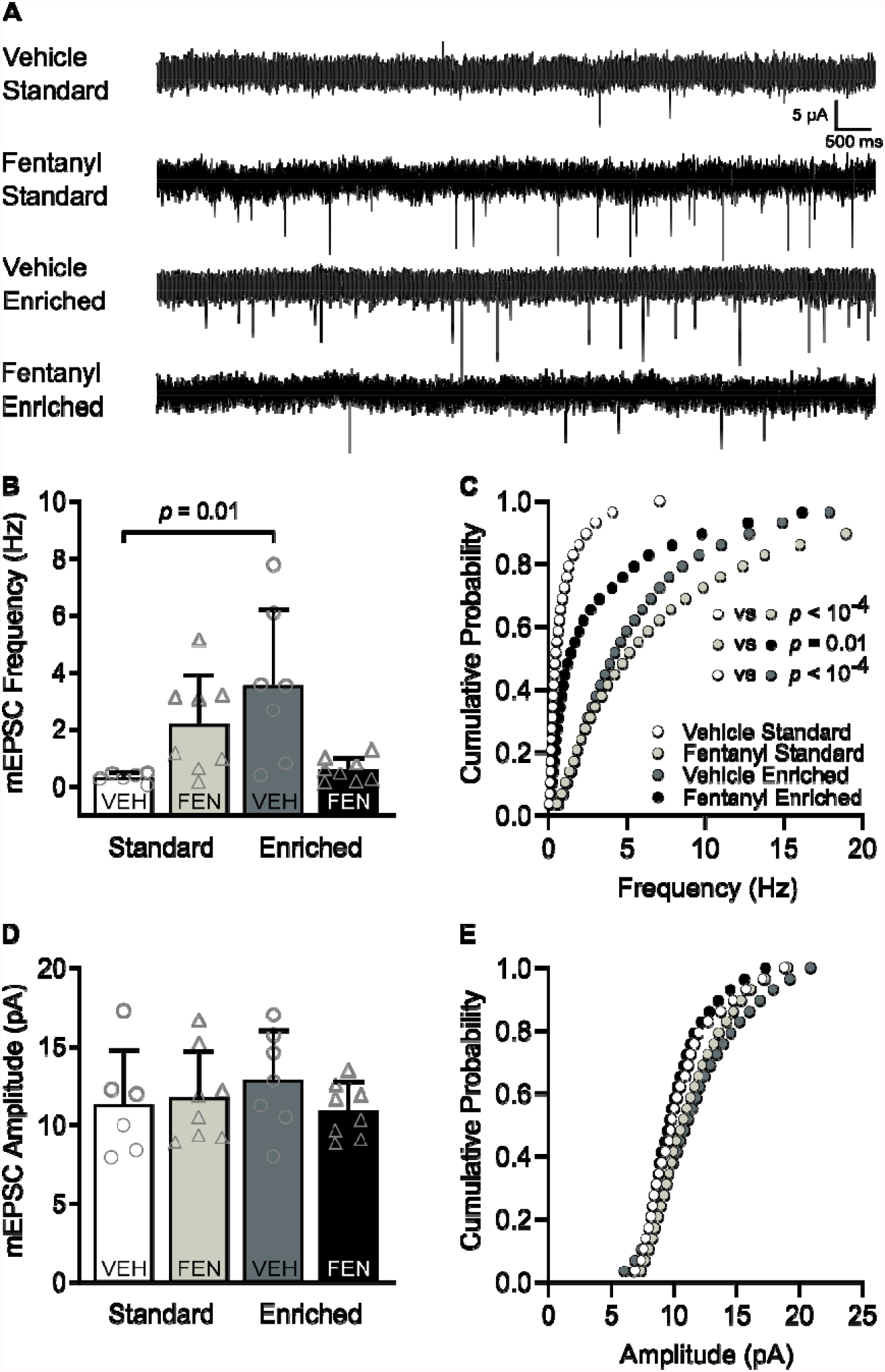
Environmental enrichment restores changes in mEPSC frequency induced by perinatal fentanyl exposure. (**A**) Representative traces of miniature excitatory postsynaptic currents (mEPSCs). (**B**) Grouped data of the averaged events from each neuron and animal indicating that mEPSC frequency is higher after environmental enrichment, compared to controls. (**C**) Cumulative probability curves of the frequency suggest perinatal fentanyl exposure results in increased mEPSC frequency, and that raising fentanyl-exposed mice with environmental enrichment reverses this increase. Enrichment also increased mEPSC frequency of vehicle control mice independent of perinatal fentanyl exposure. (**D, E**) There were no differences in mEPSC amplitudes. *n* = 6-8 mice/group, 1-2 neurons/mouse. Data depict means, and 95% confidence intervals.

Given that averaging mEPSC events can be confounded by cell-to-cell differences in vesicle release dynamics and/or postsynaptic cluster sizes (Wu et al., 2007), we further assessed these differences in mEPSC frequency by examining their cumulative probability (Fig. 6C; Kruskal-Wallis test, *H* = 42.61, *p* < 10^−4^, Pη^2^ < 0.008, *small effect*). Since we observed measurable differences among the groups in the total number and range of release events, we used a quantile-based approach described by Hanes et al., 2020 (Hanes et al., 2020) to control for these unequal distributions. Neurons from fentanyl-exposed, standard-housed mice had higher mEPSC frequency than neurons from vehicle-exposed, standard-housed mice (Dunn’s post hoc, *p* < 10^−4^, Glass’ delta = 2.98, *large effect*) and environmental enrichment normalized this increase in mEPSC frequency (Dunn’s post hoc, *p* = 0.01, Glass’ delta = 0.52, *medium-effect*).

Environmental enrichment also enhanced mEPSC frequency in vehicle-treated conditions relative to neurons from mice raised in standard housing (Dunn’s post hoc, *p* < 10^−4^, Glass’ delta = 3.34, *large effect*). In contrast, there was no significant interaction nor main effect on the average (Fig. 6D; Two-way ANOVA, *p* > 0.05) or cumulative probability (Fig. 6E; Kruskal-Wallis test, *p* > 0.05) of mEPSC amplitudes among the groups. These data suggest that both environmental enrichment and perinatal fentanyl exposure, independently establishes a higher frequency of miniature excitatory release at this synapse. Among perinatal fentanyl exposed mice, environmental enrichment restores the mEPSC frequency to a more quiescent state, resembling that of vehicle-exposed, standard-housed mice.

## Discussion

We hypothesized that environmental enrichment would mitigate the behavioral and synaptic deficits of perinatal fentanyl exposure. We administered fentanyl citrate in the drinking water of pregnant mouse dams throughout their pregnancy and until their litters were weaned at postnatal day (PD) 21. We acknowledge that opioid use among women likely occurs prior to pregnancy and contributes to both short- and long-term deleterious outcomes (Byrnes and Vassoler, 2018). To avoid effects on sexual behavior and conception (Johnson et al., 2011), we began fentanyl exposure after copulatory plugs were identified. This exposure paradigm increases litter mortality rate and decreases litter size without influencing maternal care behavior, recapitulating opioid use among pregnant women (Alipio et al., 2021b). Some studies stop opioid exposure at or a few days post parturition (Byrnes and Vassoler, 2018). We chose to continue fentanyl exposure until weaning because human cortical development at birth is equivalent to several days beyond PD 11 in mice (Clancy et al., 2001). Additionally, we wanted to avoid confounding effects due to changes in maternal care behavior, which can occur if opioid administration to the dam is discontinued following birth (Schlagal et al., 2021).

Consistent with our prediction, environmental enrichment attenuated the behavioral aberrations induced by perinatal fentanyl exposure, including hyperactivity, anxiety-like behavior, and sensory maladaptation. Enrichment also normalized short- and long-term potentiation (STP/LTP) and spontaneous excitatory synaptic transmission in primary somatosensory (S1) layer 2/3 neurons from perinatal fentanyl exposed mice. Notably, in naïve control mice, environmental enrichment results in attenuated STP/LTP and long-term depression (LTD), as well as increased spontaneous excitatory synaptic transmission.

### Behavior

We previously showed that perinatal fentanyl exposure results in spontaneous somatic withdrawal signs, lasting anxiety-like behavior, and sensory maladaptation without influencing the dam’s weight, food and liquid intake – recapitulating neonatal opioid withdrawal syndrome (Alipio et al., 2021b, 2021a). Here, we expand upon our model by showing that perinatal fentanyl exposure leads to hyperactivity, a prominent feature of attention-deficit hyperactivity disorder (Montarolo et al., 2019), which is often diagnosed in children that were exposed to opioids perinatally (Schwartz et al., 2021).

### Synaptic plasticity

Because perinatal fentanyl exposure led to impairments in sensory-related processing, we investigated changes in the efficacy of synaptic transmission in S1 layer 2/3. Plasticity in S1 layer 2/3 is associated with sensory adaptation and rapid storage of immediate sensory memories (Brecht, 2017). Since there is an immediate temporal requirement for sensory adaptation, we predicted that plasticity in this region is associated with adaptive tactile behavior. Whereas LTP was readily induced by pairing repeated stimuli with postsynaptic depolarization of S1 layer 2/3 neurons, this LTP induction protocol failed to induce LTP in neurons from fentanyl-exposed mice. In contrast, perinatal fentanyl exposure did not influence the induction of LTD in response to low frequency stimulation. These results are consistent with previous studies showing that prenatal exposure to morphine suppresses LTP in the dentate gyrus of juvenile rats (Niu et al., 2009; Ahmadalipour et al., 2018).

Here, we find that environmental enrichment increased the short-term potentiation of EPSCs from fentanyl-exposed mice. The potentiation was sustained in a subset of neurons up to 30 min post stimulation. We recorded out to 30 min post stimulation, which is within the accepted parameters of conventional LTP (Nicoll, 2017), to avoid possible confounding effects of a sustained timecourse (Abbas et al., 2015; Baltaci et al., 2019). We did not assess short-term depression since the resulting EPSC amplitude depression was sustained at least to 30 min post stimulation. Further studies are needed to investigate late phases of plasticity under these conditions.

### Synaptic activity

We have previously reported that perinatal fentanyl exposure suppresses evoked and spontaneous glutamate release onto S1 layer 5 neurons, and enhances glutamate release onto anterior cingulate cortex (ACC) layer 5 neurons (Alipio et al., 2021b). Here we find that fentanyl exposure enhances spontaneous glutamate release onto S1 layer 2/3 neurons. That neurons in different cortical layers and areas express differences in the lasting effects of perinatal fentanyl exposure likely reflects differences in cortical developmental trajectory. Our data suggest that neurons that develop earlier (S1 layer 2/3 neurons) express a lasting increase in excitatory synaptic activity, whereas later developing neurons (S1 and ACC layer 5 neurons) exhibit decreased synaptic activity.

The different effects on spontaneous versus evoked glutamate release is not surprising. Mechanisms that regulate these two phenomena are well documented (Ramirez and Kavalali, 2011; Kavalali, 2015). For example, brain-derived neurotrophic factor (BDNF) enhances mEPSCs but not evoked EPSCs in immature cortical neurons (Taniguchi et al., 2000). We have previously found that perinatal fentanyl exposure decreases mRNA expression of TrkB, a receptor for BDNF (Alipio et al., 2021b). Given the differential expression patterns of TrkB across cortical layers (Cabelli et al., 1996), it is possible that our fentanyl exposure paradigm interferes with the developmental trajectory of TrkB expression and subsequently impairs cortical circuit function.

### Environmental enrichment effects on behavior

Environmental enrichment restored the hyperactivity induced by perinatal fentanyl exposure, as well as the increase in anxiety-like behavior in a novel open field environment. This is consistent with studies demonstrating that rearing rodents in an enriched environment improves maladaptive phenotypic changes in preclinical models of ADHD and anxiety (Botanas et al., 2016; Korkhin et al., 2020; Yazdanfar et al., 2021). Environmental enrichment also restored the sensory maladaptation induced by perinatal fentanyl exposure. These findings might be relevant to addressing the sensory-related deficits in children that were perinatally exposed to opioids.

Other preclinical studies have shown that environmental enrichment can ameliorate behavioral changes induced by perinatal exposure to other drugs of abuse, including nicotine, cocaine, morphine, ethanol, antiadrenergic, and antihypertensive drugs (Ryan and Pappas, 1990; Dow-Edwards et al., 2014; Mychasiuk et al., 2014; Ahmadalipour et al., 2018; Wille-Bille et al., 2020; Yazdanfar et al., 2021). These results suggest that enrichment may benefit infants and children with such exposures. Importantly, our data also support that environmental enrichment may be a favorable intervention during early development, since it does not produce untoward behavioral outcomes on its own, at least not in the behavioral outcomes assessed in the current study.

### Environmental enrichment effects on synaptic plasticity

Environmental enrichment restored the perinatal fentanyl exposure-induced impairment in LTP in S1 layer 2/3 neurons. This is consistent with findings that environmental enrichment restores stress-induced reduction of LTP in hippocampal dentate gyrus and prefrontal cortical neurons, and that enrichment does not further increase LTP in non-stressed controls (Wang et al., 2020; Wu and Mitra, 2020).

We also found that environmental enrichment suppressed the ability of S1 layer 2/3 neurons to express either LTP or LTD. These findings are consistent with previous reports showing that environmental enrichment-induced plasticity occludes further potentiation of S1 layer 2/3 and 4 neurons (Mégevand et al., 2009). Similarly, enrichment reduces or blocks the induction of LTP in the dentate gyrus (Feng et al., 2001; Irvine et al., 2006; Eckert et al., 2010), perhaps by occluding LTP (Foster et al., 1996). The occlusion of experimentally driven plasticity by sensory experience is consistent with findings that cortical neurons may reach a modification ceiling following training on a sensorimotor task (Rioult-Pedotti et al., 2000). However, despite occlusion of LTP by natural behavior, further potentiation remains possible through other synaptic strengthening mechanisms (Clem et al., 2008). It remains to be determined if further plasticity can be induced under the conditions here, perhaps by way of different induction protocols.

### Environmental enrichment effects on synaptic activity

Environmental enrichment restored the amplified spontaneous excitatory synaptic transmission in S1 layer 2/3 neurons induced by perinatal fentanyl exposure, and increased it in control mice. The contrasting effects of enrichment on fentanyl and on control mice reflect the multifactorial mechanisms in which enrichment influences cortical synaptic transmission (Baroncelli et al., 2010). Relevant findings demonstrate that environmental enrichment restores the decreased mEPSC frequency in S1 layer 2/3 neurons induced by sensory deprivation, and increases mEPSC frequency in control, non-deprived mice (Zheng et al., 2014). The mechanisms involved in the increased excitatory synaptic activity induced by perinatal fentanyl exposure and the restoration by environmental enrichment has yet to be determined.

Converging evidence from longitudinal human studies and preclinical models demonstrate that perinatal exposure to opioids results in long-lasting molecular, circuit, network, and behavioral aberrations. Current treatment options are aimed at relieving acute symptoms exhibited by newborns with neonatal opioid withdrawal. No such treatments are available for the developing child or adolescent. Our results provide insights into the underlying circuit changes involved in the lasting anxiety and sensory-related deficits induced by perinatal fentanyl exposure, and suggest that environmental enrichment may be leveraged to ameliorate or reverse the lasting deleterious effects of this early opioid exposure.

## Acknowledgements

JBA and LMR conceptualized the project, designed and performed the experiments, analyzed and interpreted the data, and co-wrote the manuscript; MP assisted in data analysis; AK supervised the research, performed analyses, and co-wrote the manuscript. We thank Urja Kuppa for preliminary analysis of miniature excitatory postsynaptic currents.

